# An Updated Structure of Oxybutynin Hydrochloride

**DOI:** 10.1101/2024.06.05.597682

**Authors:** Jieye Lin, Guanhong Bu, Johan Unge, Tamir Gonen

## Abstract

Oxybutynin (Ditropan), a widely distributed muscarinic antagonist for treating the overactive bladder, has been awaiting a definitive crystal structure for nearly 50 years due to the sample and technique limitations. Past reports used powder X-ray diffraction (PCRD) to shed light on the possible packing of the molecule however a 3D structure remained elusive. Here we used Microcrystal Electron Diffraction (MicroED) to successfully unveil the 3D structure of oxybutynin hydrochloride. We identify several inconsistencies between the reported PXRD analyses and the experimental structure. Using the improved model, molecular docking was applied to investigate the binding mechanism between M_3_ muscarinic receptor (M_3_R) and (*R*)-oxybutynin, revealing essential contacts/residues and conformational changes within the protein pocket. A possible universal conformation was proposed for M_3_R antagonists, which is valuable for future drug development and optimization. This study underscores the immense potential of MicroED as a complementary technique for elucidating the unknown pharmaceutical crystal structures, as well as for the protein-drug interactions.

---

Oxybutynin, marketed as “Ditropan”, is a muscarinic antagonist for overactive bladder treatment. It was first approved for medical use in the United States in 1975 (nearly 50 years ago).^1-3^ Unlike mirabegron (Myrbetriq), a β_3_ adrenoceptor agonist responsible for bladder relaxation,^4^ oxybutynin targets the M_3_ muscarinic receptor (M_3_R) as an antagonist, effectively suppressing bladder contraction by preventing the binding of acetylcholine^5,6^ and conformational change in M_3_R needed for downstream G_q_/_11_ signaling.^7^ The commercial formulation of oxybutynin is a racemic mixture which contains *R*- and *S*-enantiomers. The molecule features a phenyl ring, a cyclohexyl ring, a hydroxyl group, and an ester-linked aliphatic chain with a carbon-carbon triple bond (C≡C) and a dimethylamine group, all connected via a chiral carbon atom (Figure 1A). Traditionally, the single-crystal X-ray diffraction (SC-XRD) is the primary technique for elucidating 3D crystal structure of M_3_R antagonists like tiotropium (CSD entry: GUYGOX, 2010),^8^ trospium (CSD entry: IPILUQ, 2016),^9^ solifenacin (CSD entry: URATAK, 2016),^10^ *etc*. However, SC-XRD was not suitable for oxybutynin hydrochloride,^11^ because this drug typically forms micro or nano sized crystals in a seemingly amorphous powder that is not amenable to structure determination using this approach. The powder X-ray diffraction (PXRD) was used as an alternative. However, the PXRD structure of oxybutynin hydrochloride hemihydrate was controversial because it was inconsistent with the 2D chemical structure, for example with the absence of an O atom in the ester bond.^13^ Previous PXRD studies revealed photoreduction of the C≡C bond in oxybutynin when exposed to either synchrotron or in-house X-ray sources^13^ and refinement of radiation-damaged data led to inaccuracies in structural coordinates. PXRD may sometimes also encounter problems like line broadening, peak overlapping, and time-consuming computer based calculation^12,13^ for complex samples. Other techniques, like solid-state NMR, have not been reported for oxybutynin.^14^ Due to the various problems as outlined above, the crystal structure of oxybutynin hydrochloride, which is the anhydrous form used in the pharmaceutical formulations, has been undetermined for nearly 50 years. Nonetheless, it has widely been prescribed, ranking as #102 most prescribed medicines in the United States in 2021, with ∼7 million prescriptions.^15^

**Figure 1.**
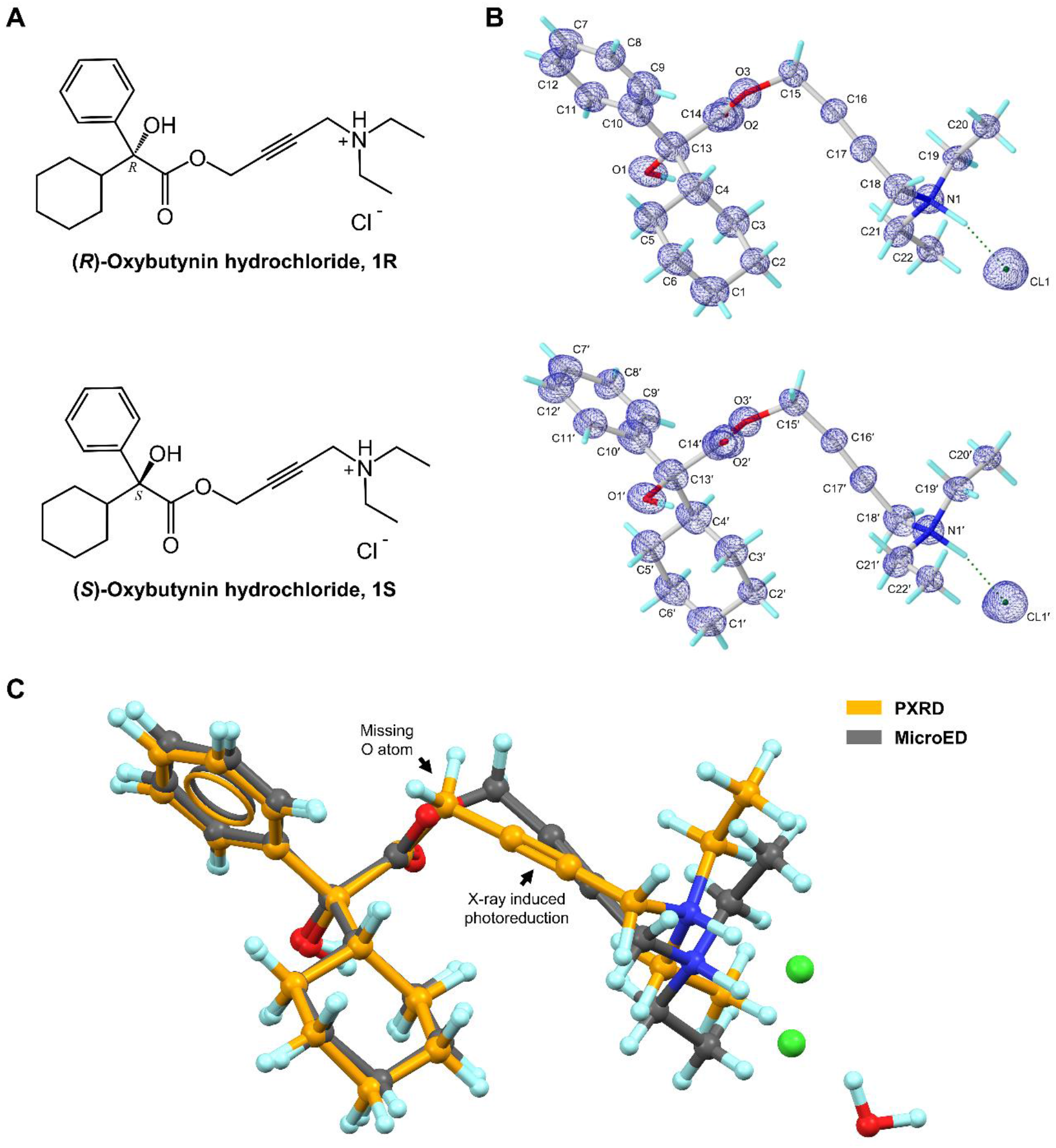
Chemical (A) and MicroED structures (B) of (*R*)-oxybutynin hydrochloride **1R** and (*S*)-oxybutynin hydrochloride **1S**. 2F_o_-F_c_ density map was shown in blue mesh. (C) Overlay of MicroED and literature reported PXRD structure of **1R** showing the missing O atom error and X-ray induced photoreduction in C≡C bond in the PXRD structure. MicroED structure was colored in grey, PXRD structure was colored in orange.

The development of microcrystal electron diffraction (MicroED) technique bypasses the crystal size limitations of X-ray diffraction.^16,17^ Due to the larger atomic cross-section and increased elastic scattering with matter, MicroED is especially suitable for micro- or nano-sized crystals, requiring crystals with only a billionth of the size needed for SC-XRD.^18^ MicroED data collection is conducted at cryogenic temperature and using high vacuum in a cryo-TEM, employing an ultralow radiation dose rate (∼0.01 e^-^/Å^2^/s) during the fast continuous-rotation data collection (∼1 minute exposure per crystal),^19^ which significantly reduces the radiation damage. The growing application of MicroED to pharmaceutical molecules has unveiled elusive crystal structures of drugs that have been in medical use for decades, for example, paritaprevir,^20,21^ simeprevir,^21^ indomethacin,^22^ meclizine,^23^ *etc*. The new structural insights obtained not only serve as a complement to existing literature but are also crucial for drug development. In this study, MicroED was applied to successful crystal structure determination of oxybutynin hydrochloride after nearly 50 years of medical use. The sub-atomic structure was directly solved from a ∼1 µm sized crystal (Figure S1, Supporting Information) using ultralow electron doses. The statistics of the data collection parameters showed no signs of radiation damage, enabling the revision of and significant enhancement over the former PXRD structure.

As oxybutynin is an M_3_R antagonist, it inhibits the entry of biological agonist into M_3_R and the required conformational change of M_3_R for downstream G_q_/_11_ signaling.^5-7^ Previous studies revealed the antimuscarinic activity of oxybutynin stereo-selectively dependent on its (*R*)-enantiomer, because of the distribution, binding difference, and other factors in human plasma.^24-26^ There is no experimental report about the structure between M_3_R and oxybutynin in complex, likely due to the challenges of handling this membrane protein *in vitro*. Using the fusion protein M_3_R with a T4 lysozyme, the complex structures like M_3_R/tiotropium (PDB entry: 4U15) and M_3_R/*N*-methyl scopolamine (PDB entry: 4U16) have been successfully determined.^6^ On this basis, we employed molecular docking to analyze the binding between M_3_R and (*R*)-oxybutynin, uncovering (1) the essential contacts in the protein pocket; (2) the conformational changes of (*R*)-oxybutynin from the drug-formulation state to the biologically active state. Comparison of the predicted complex structure with three other M_3_R antagonists highlights the universal binding geometry and residues necessary for function. Structural insights navigate the future drug development and optimization.

The MicroED sample preparation of oxybutynin hydrochloride **1** followed the procedure described in the literature (See details in Supporting Information).^18^ The continuous carbon-coated grid containing the crystals was inserted to a 200 kV (0.0251 Å wavelength) Talos Arctica Cryo-TEM (Thermo Fisher) equipped with a CetaD CMOS camera and EPUD software.^27^ The thickness of crystals is crucial for the quality of MicroED data, therefore only the thinner crystals with the suitable visual contrast (Figure S1A, Supporting Information) were selected under the imaging mode (SA 3400x). Those selected crystals were calibrated to eucentric heights in order to steadily maintain them within the beam during the continuous rotation. The MicroED data was collected under the diffraction mode (741 mm diffraction length) using the parallel beam settings (0.0098 e^-1^/Å^2^/s). Typical data collection used 0.5 s exposure time per frame, and a constant rate of 2° per second over the wedge of 130° (-65° to +65°), ensuring the ultralow dose (0.65 e^-^/Å^2^) for each dataset (see Supporting Information for more details). The optimal MicroED dataset (mrc format) was converted to images (smv format) using mrc2smv software (https://cryoem.ucla.edu/microed).^27^ The converted frames were indexed, integrated, and scaled in XDS,^28,29^ achieving an overall completeness of 83.9%. Intensities were converted to SHELX hkl format using XDSCONV^29^ and *ab initio* solved by SHELXT^30^ at the resolution of 0.87 Å in a centrosymmetric monoclinic C2/c space group with the unit cell of **a**=38.85 Å, **b**=7.92 Å, **c**=14.43 Å, **α**=90.000**°**, **β**=110.560°, **γ**=90.000°. The SHELXL refinement yielded a final R_1_ value of 18.42% (Table S1, Supporting Information).^31^ The positions of non-hydrogen atoms were accurately determined from the sub-atomic charge density map (Figure 1B). The polar H atoms were located in the difference map, while the non-polar H atoms were placed using riding models. Comparing the back-calculated PXRD pattern of **1** and the literature-reported oxybutynin hydrochloride hemihydrate indicated different crystalline forms, *i*.*e*. anhydrous and hydrate forms (Figure S2, Supporting Information). Direct comparison of their unit cell parameters showed 1.7 Å expansion in the longest axis, and 264 Å^3^ expansion in cell volume.^13^ Compared with the reported PXRD structure, the data collected here resolved the missing oxygen atom in the ester bond and showed the normal C≡C bond without any photoreduction, and is now included in the MicroED model (Figure 1C).^13^

Both two enantiomers, (*R*)-oxybutynin hydrochloride **1R** and (*S*)-oxybutynin hydrochloride **1S** were identified within the unit cell, differed by configurations at the chiral center (C13/C13′ atoms, see chemical notations in Figure 1B). **1R** and **1S** densely packed as repeated layers (Figure 2A; Figure S3, Supporting Information). Each layer was formed by the stacking (along *a*-axis) and direct translation (along *b*- and *c*-axes) of four **1R**/**1S** molecules as a repeating unit. The elongation of crystal lattice is a result from six chloride anion-mediated hydrogen bonds along three axes, for example, the N1−H**…**Cl1/N1′−H…Cl1′ hydrogen bonds along *c*-axis (3.05 Å); the O1−H…Cl1′/O1′−H…Cl1 (3.09 Å) and C19−H…Cl1′/C19′−H…Cl1 (3.59 Å) hydrogen bonds along *a*- and *b*- axes (Figure 2B; Table S2, Supporting Information). Chloride anions are hydrogen bonding acceptors posed between **1R** and **1S**, which adjust the lipophilicity and bioavailability of the drug, *e*.*g*. the predicted *n*-octanol/water partition coefficient logP_o/w_ lowered down to 1.50 from the base form 3.74.^32^ Removal of this anions makes the drug more lipophilic and better fit with the hydrophobic environment when interacting with its muscarinic receptor. Medium T-shaped π-stacking interactions (4.93 Å) are found in **1R** and **1S** layers along *c*-axis which strengthen the packing but do not interlink to extend the crystal packing (Figure 2C).

**Figure 2.**
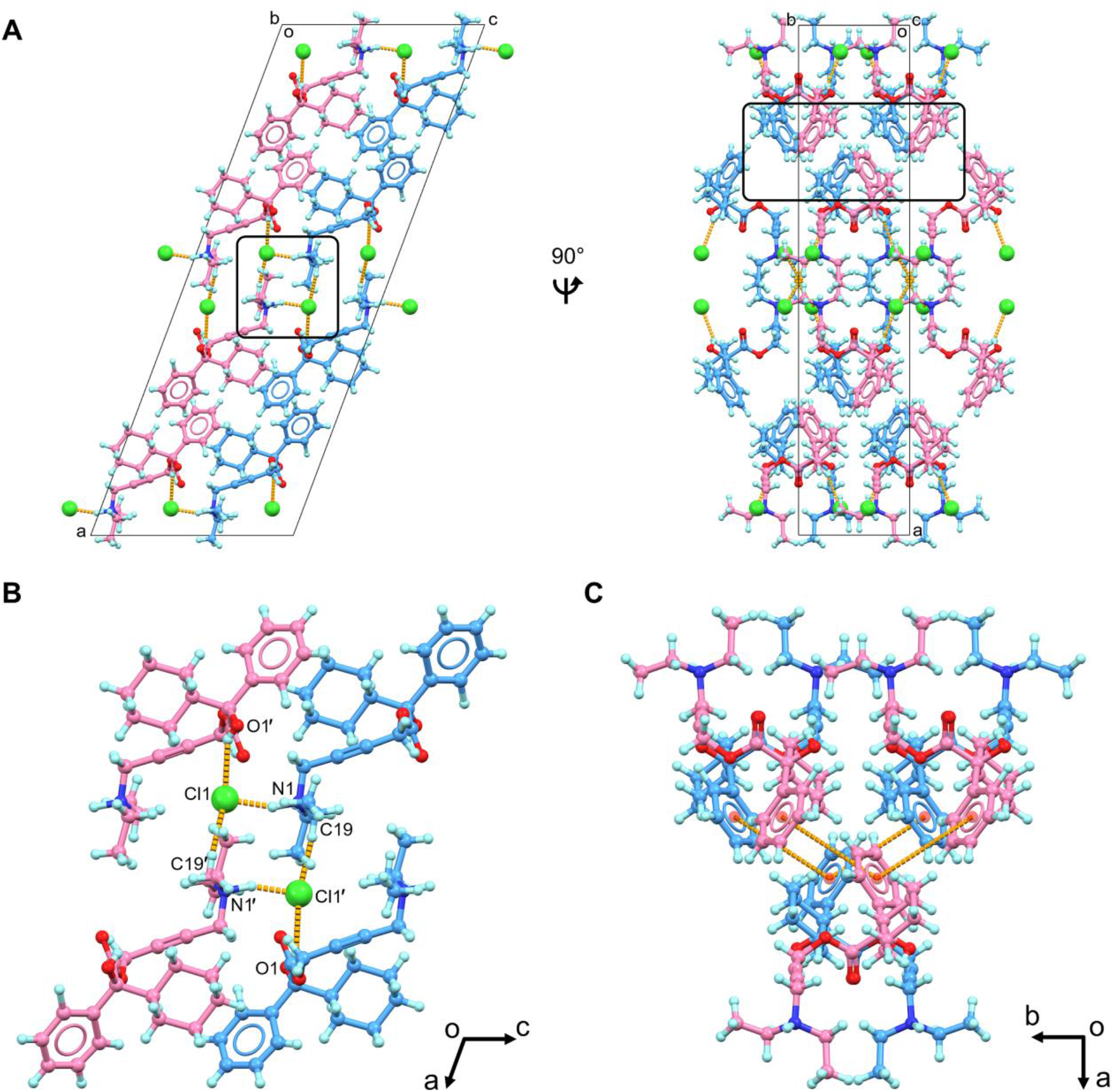
(A) Packing diagram of oxybutynin hydrochloride **1**, viewed along *b* and *c* axes; (B) Hydrogen bonding interactions in **1**, viewed along *b* axis; (C) π-stacking interactions in **1** (less than 5 Å). **1R** was colored in blue, **1S** was colored in violet. Hydrogen bonding and π-stacking interactions were represented by the dashed lines in orange. Cl^**−**^ anions were highlighted in spacefill style. Extra Cl^**−**^ anions were omitted in Figures 2B and 2C for clarification.

Each **1R**/**1S** enantiomer contains a chiral center at C13/C13′ atom, which is connected by one hydroxyl group, one phenyl ring, one cyclohexyl ring, and an ester-bonded linear aliphatic chain containing a triple C≡C bond and a diethylamine group (Figure 1B). Examining the structural parameters in **1R** and **1S** revealed no apparent distortions in bond lengths, and negative density is not observed in the difference map, indicating no radiation damage occurred during the experiment. For example, the triple C≡C bond is 1.23 Å, contrary to the prior observation of X-ray induced photoreduction of triple bond.^13^ Most of bond angles in **1R**/**1S** suggest a sp^3^ geometry, for example, the average C−C−O/C bond angles around C13/C13′ atom and C−N−C/H bond angles around N1/N1′ atom have the average value of ∼109°. Although the incorporation of ester group (C14/C14′, O2/O2′, O3/O3′ atoms) and triple C≡C bond (C16/C16′, C17/C17′ atoms) restrict the conformational flexibility of **1R/1S**, there are at least **9** torsion angles (C−C, C−O and C−N bonds) that theoretically have high rotational freedom (Figure S4, Supporting Information). Most of them adopt the *staggered* conformer in the drug formulation state, while they can be altered to less energetically favorable conformation upon binding in the protein pocket. Structures in the drug formulation state and the biologically active state were investigated below (Figure S4, Supporting Information).

Previous studies indicated the antimuscarinic activity of **1** to M_3_R stereo-selectively resides by **1R** because of the distribution, binding difference, and other factors in human plasma.^1,24-27^ To date, no experimental protein-ligand complex structure of M_3_R and **1R** has been reported to show the exact binding mechanism. The challenges arose from the purification, stabilization, and structure determination of M_3_R, a member of the difficult G protein-coupled receptors. Computational approaches such as molecular docking on the other hand has increasingly become a powerful tool in the prediction of protein-drug binding complexes, effectively addressing the questions involved. Herein, this method was employed to predict the binding of **1R** in M_3_R. The prior research has elucidated the complex structures of a fusion protein of M_3_R with T4 lysozyme (denoted below as “M_3_R” for clarification) binding with antagonists like tiotropium (PDB entry: 4U15) and *N*-methyl scopolamine (PDB entry: 4U16).^6^ Given the chemical similarities between **1R** and these two antagonists, the fusion protein (PDB entry: 4U15) was used as a rigid receptor in molecular docking. The MicroED structure of **1R** was extracted and served as the flexible ligand in molecular docking. The AutoDock Vina^33,34^ setup followed the procedures described in the Supporting Information and the complex structure was analyzed by Protein-Ligand Interaction Profiler (PLIP).^35^

**1R** was docked into a hydrophobic pocket of M3R, with the geometry observed in the M_3_R/**1R** complex similar to that of M_3_R/tiotropium (PDB entry: 4U15). The diethylamine group in **1R** engages in hydrophobic interactions with a tyrosine lid (Tyr148, Tyr506, Tyr529, Tyr 533), and contacts with Asp147 and Ser151 residues via one salt bridge and one hydrogen bond, respectively (Figure 3B; Table S3, Supporting Information). The carbonyl group and hydroxyl group of **1R** are hydrogen bonded with the amide group and the carbonyl group of Asn507 side chain at 2.94 Å and 2.90 Å, respectively. The latter contact causes the hydroxyl group to shift from a *staggered* conformation in the drug formulation state to an *eclipsed* conformation in the biological state. The terminal phenyl and cyclohexyl rings on the other side of **1R** are involved in seven hydrophobic interactions with the aliphatic carbon atoms of Asn152, Val155, Trp199, Thr231, Thr234, Ala235 and Ala238 residues, possibly influencing the conformational changes in M_3_R. The occupancy of **1R** in M_3_R blocks the entry of biologically active agonist.

**Figure 3.**
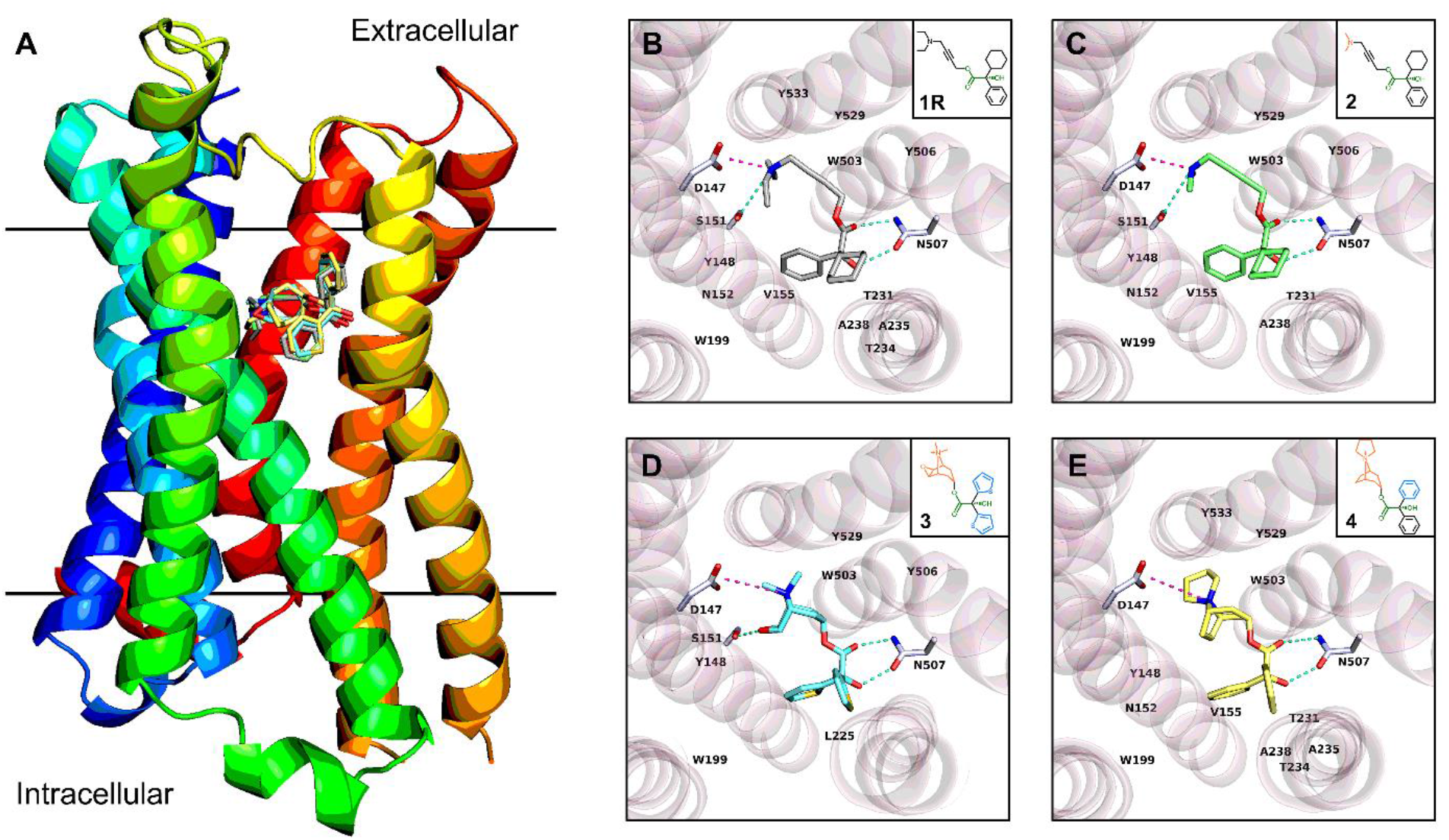
(A) Overlay of protein-drug interaction diagram of complexes between M_3_R and four antagonists; (B) Topside view of M_3_R/**1R** complex structure predicted by molecular docking; (C) Topside view of M_3_R/**2** complex structure predicted by molecular docking; (D) Topside view of M_3_R/**3** complex structure determined by X-ray diffraction (PDB entry: 4U15); (E) Topside view of M_3_R/**4** complex structure predicted by molecular docking. Hydrogen bonding interactions were colored by the dashed line in greencyan, and salt bridges were colored by the dashed line in light magenta. π-stacking, π-cation and hydrophobic interactions were omitted for clarification (see details in Tables S3-S6, Supporting Information). The fusion parts of T4 lysozyme were omitted for clarification. Compounds were symbolled as **1**-**4**: (*R*)-Oxybutynin **1R**, (*R*)-4-Dimethylamino-2-butynyl-2-cyclohexyl-2-hydroxy-2-phenylacetate hydrochloride **2**, Tiotropium **3**, Trospium **4**.

While interacting with the protein target, the ligand is not necessarily in its energetically minimal state, since the entropy cost is compensated by diverse interactions in the protein pocket.^36^ It is also applicable for **1R**, which undergoes three major conformational changes during the transition from the drug-formulation state to the biologically active state (Figure S4, Supporting Information). For example, the O3−C15 bond has a ∼230° rotation from a conformation between the *staggered* and *eclipsed* conformer (-90°) to an *eclipsed-like* conformer (140°). This less favorable conformation is compensated by the interactions with Tyr506 and Tyr529 residues. The linear geometry from C15 to C18 atoms is maintained by the C≡C triple bond, while the large rotation of C17−C18 bond from -129° to 144° reoriented the terminal diethylamine group. The above conformational changes together with the rotation of N1−C21 and N1−C19 bond positioned the ethyl parts towards the Asp147, Ser151, Tyr148 and Tyr533 residues, facilitating more salt bridges, hydrogen bonds, and hydrophobic interactions (Figure 3B; Table S3, Supporting Information). Such conformation cannot be easily achieved by **1S** which partially explains the stereo-selectivity, for example, the carbonyl O2′ atom in the opposite direction can hardly be involved in the hydrogen bonding interaction with Asn507, the rotation of dimethylamine group is therefore more restricted than **1R**.

To validate whether the above geometry is a universal pose among M_3_R antagonists and to figure out the necessary residues involved in binding, three M_3_R antagonists **2**-**4** were selected: (*R*)-4-dimethylamino-2-butynyl-2-cyclohexyl-2-hydroxy-2-phenylacetate hydrochloride **2**, tiotropium **3** and trospium **4**,^6,9,37^ which have analogous chemical structures and functional groups with **1R** (Figures 3B-3E). Four compounds differ by their amine groups, for example, the diethylamine group in **1R**, the dimethylamine group in **2**, and the bicyclic rings containing a quaternary amine in **3** and **4** (Figures 3B-3E). The complex structures between M_3_R and **1R, 2** and **4** were calculated (see Supporting Information for more details), while the experimentally solved complex structure between M_3_R and **3** was retrieved from PDB (PDB ID 4U15).^6^ All four complex structures exhibited a comparable protein-drug binding pose (Figure 3; Tables S3-S6, Supporting Information). For example, a salt bridge is formed between Asn147 residue and amines in **1**-**4**, which is further stabilized by a hydrogen bond with a Ser151 residue; multiple hydrophobic/π-cations interactions with Tyr148, Trp503 and Tyr529 residues. The carbonyl and hydroxyl groups generate two hydrogen bonds with Asn507. While different rings and their rotations led to different hydrophobic contacts in this pose, for instance, the phenyl and cyclohexyl rings in **1R, 2** and **4** are larger and more lipophobic, which are contacted by aliphatic carbon atoms of Asn152, Val155, Trp199, Thr231 and Ala238 residues, whereas the thiophenyl group in **3** predominantly interacts with Trp199 and Leu225 residues.

In this study, we utilized MicroED to successfully determine the elusive crystal structure of oxybutynin hydrochloride, a compound that has been in widespread medical use for nearly 50 years. The result is particularly noteworthy because it: (1) enables a radiation damage free model by using cryogenic conditions and very low electron doses.^19^ to (2) improve the previous PXRD structure (*i*.*e*. the missing oxygen atom in the ester bond); Based on the crystal structure of oxybutynin in its drug formulation state, we conducted molecular docking to analyze the binding mechanism with M_3_R. This analysis identifies crucial M_3_R residues for interactions, pinpoints the conformational changes from the drug-formulation state to the biologically active state, and proposes a possible universal conformation for M_3_R antagonists, which are supportive for advancing and optimizing the next-generation M_3_R antagonists (*e*.*g*. conformational constraints). This study underscores the immense potential of MicroED as a complementary technique for elucidating the unknown pharmaceutical crystal structures which have previously been hindered by issues like crystal size, chemical/physical properties, and radiation damage *etc*. and were unachievable by existing techniques. By combining MicroED with computational techniques like molecular docking, protein-drug interactions can be extensively investigated.

## Acknowledgements

The authors thank Michael W. Martynowycz for support and discussions. This study was supported by the National Institutes of Health P41GM136508. Portions of this research or manuscript completion were developed with funding from the Department of Defense MCDC-2202-002. Effort sponsored by the U.S. Government under Other Transaction number W15QKN-16-9-1002 between the MCDC, and the Government. The US Government is authorized to reproduce and distribute reprints for Governmental purposes, notwithstanding any copyright notation thereon. The views and conclusions contained herein are those of the authors and should not be interpreted as necessarily representing the official policies or endorsements, either expressed or implied, of the U.S. Government. The PAH shall flow down these requirements to its sub awardees, at all tiers. The Gonen laboratory is supported by funds from the Howard Hughes Medical Institute.

