## Supporting Information for "An Updated Structure of Oxybutynin Hydrochloride"

### Methods

#### Materials.

Oxybutynin hydrochloride, 4-(diethylamino)but-2-ynyl (*R*/*S*)-2-cyclohexyl-2-hydroxy-2-phenyl-acetate hydrochloride, was commercially purchased from InvivoChem and used as received, without further recrystallization.

#### Grid preparation.

The sample preparation followed the procedure as described previously.[1] One carbon-coated copper grid (200-mesh, 3.05 mm O.D., Ted Pella Inc.) was pretreated with 15 mA negative glow-discharge plasma for 60 s using PELCO easiGlow (Ted Pella Inc.). Less than 1 mg of powdery compounds was mixed with the grid in a 10 mL scintillation vial with 30 s gentle shaking. The grid was clipped outside using c-ring and autogrid clip at room temperature.

#### MicroED data collection.

The clipped grid was loaded in Thermo Fisher Talos Arctica Cryo-TEM (200 kV, ~0.0251 Å), which is equipped a CMOS CetaD camera (4096 × 4096 pixels) and EPUD software (Thermo Fisher).[2] Screening of microcrystals was done in the imaging mode (SA 3400×). Only the thin crystals with a certain brightness contrast were selected. Their eucentric heights were calibrated to maintain the crystals inside the beam during the continuous rotation. The MicroED data was collected in the diffraction mode at the camera length of 741 mm (the calibrated sample-detector distance) and microprobe beam size 11 under the parallel beam condition (45.2% C2 intensity). A 70 µm C2 aperture and a 50 µm selected area (SA) aperture were used to result in an approximately 1.4 µm width beam area (0.0098 $e^{-1}/Å^2/s$).[3] Typical data collection used a constant rotation rate of 2° per second over an angular wedge of 130° from -65° to +65°, with 0.5s exposure time per frame, resulting a total dose of 0.65 $e^-/Å^2$ for each dataset.

**MicroED data processing.**

MicroED data was saved in mrc format and converted to smv format using the mrc2smv software (https://cryoem.ucla.edu/microed).[2] The converted frames were indexed and integrated by XDS.[4,5] One dataset with the highest resolution (0.87 Å) was scaled using XSCALE to achieve 83.9% completeness.[5] Intensities were converted to SHELX hkl format using XDSCONV[5] and *ab initio* solved by SHELXT.[6] The structure was refined by SHELXL[7] using Shelxle[8] as a graphical interference to yield the final MicroED structure (Table S1).

**Molecular Docking**

The ligand structures from MicroED structure **1R** and X-ray structure **2** (CSD entry: MBCHPA) [9] and **4** (CSD entry: IPILUQ)[10] were extracted. The amine hydrogens were removed based on pKa value.[11] All the water and $Cl^-$ anions were removed. The ligand structures were imported to AutoDock Tools 1.5.7,[12] making all active torsion bonds rotatable (Figure S4). The protein structure was downloaded from PDB (Protein Data Bank, https://www.rcsb.org/) entry 4U15.[13] Ligands, ions, water were removed using Pymol 2.5.5.[14] The resulting protein structure was modified by adding hydrogen atoms and charges as computed by AutoDock Tools 1.5.7.[12]

The protein cavity was analyzed by CB-Dock2 webtool[15] at the center of x,y,z = (17, 85, 56), which is consistent with the orthosteric binding site observed in PDB structure 4U15.[13] A grid box measuring 18.75 Å × 18.75 Å × 18.75 Å with 0.375 Å spacing was positioned. During the docking in AutoDock Vina 1.1.2,[16,17] the ligands were set to be flexible (Figure S4), while the protein was treated with rigid model. The docked complex with minimum binding energy was analyzed in Protein-Ligand Interaction Profiler (PLIP) web tool and Pymol 2.5.5 (Figure 3 and Tables S3-S6).[14,18]

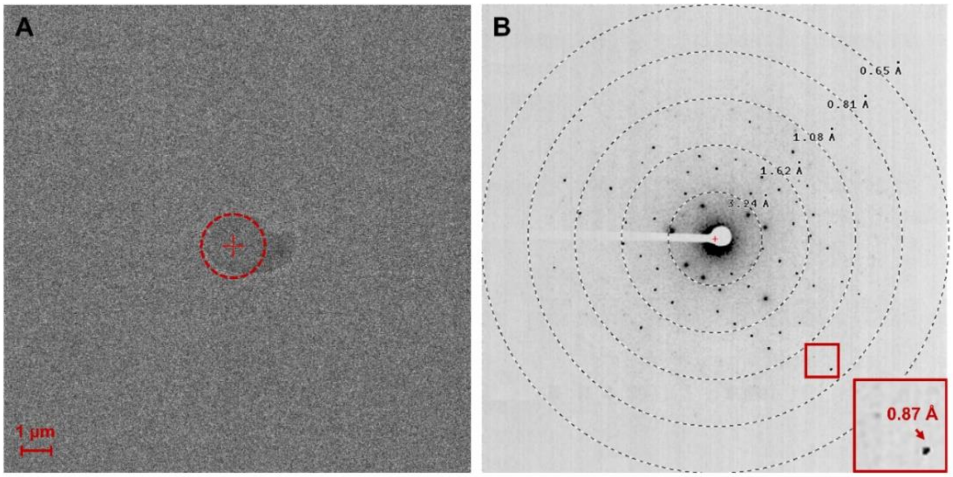


**Figure S1** Crystal appearance and diffraction pattern under the TEM. (A) Image of oxybutynin hydrochloride **1** under the imaging mode (SA 3400×). The diffraction beam area was highlighted in dashed red circles; (B) Diffraction pattern of oxybutynin hydrochloride **1** under diffraction mode (741 mm).

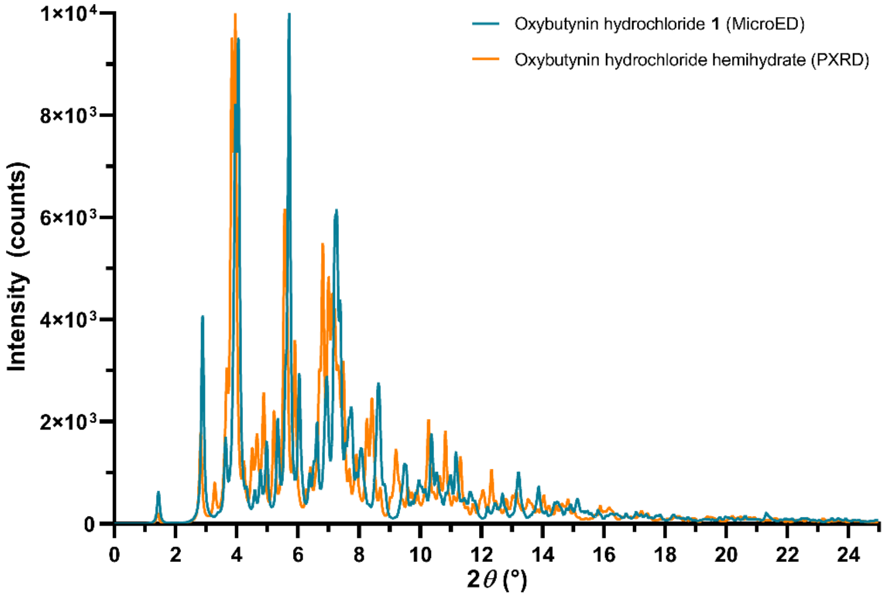


**Figure S2** Overlay of PXRD patterns of oxybutynin hydrochloride **1** and the literature reported oxybutynin hydrochloride hemihydrate.[19] The PXRD patterns were back-calculated using the MicroED and PXRD structures in Mercury software[20] (wavelength was set as 0.457667Å with 0.001° 2$\theta$ step to be consistent with literature[19]), colored in green and orange, respectively.

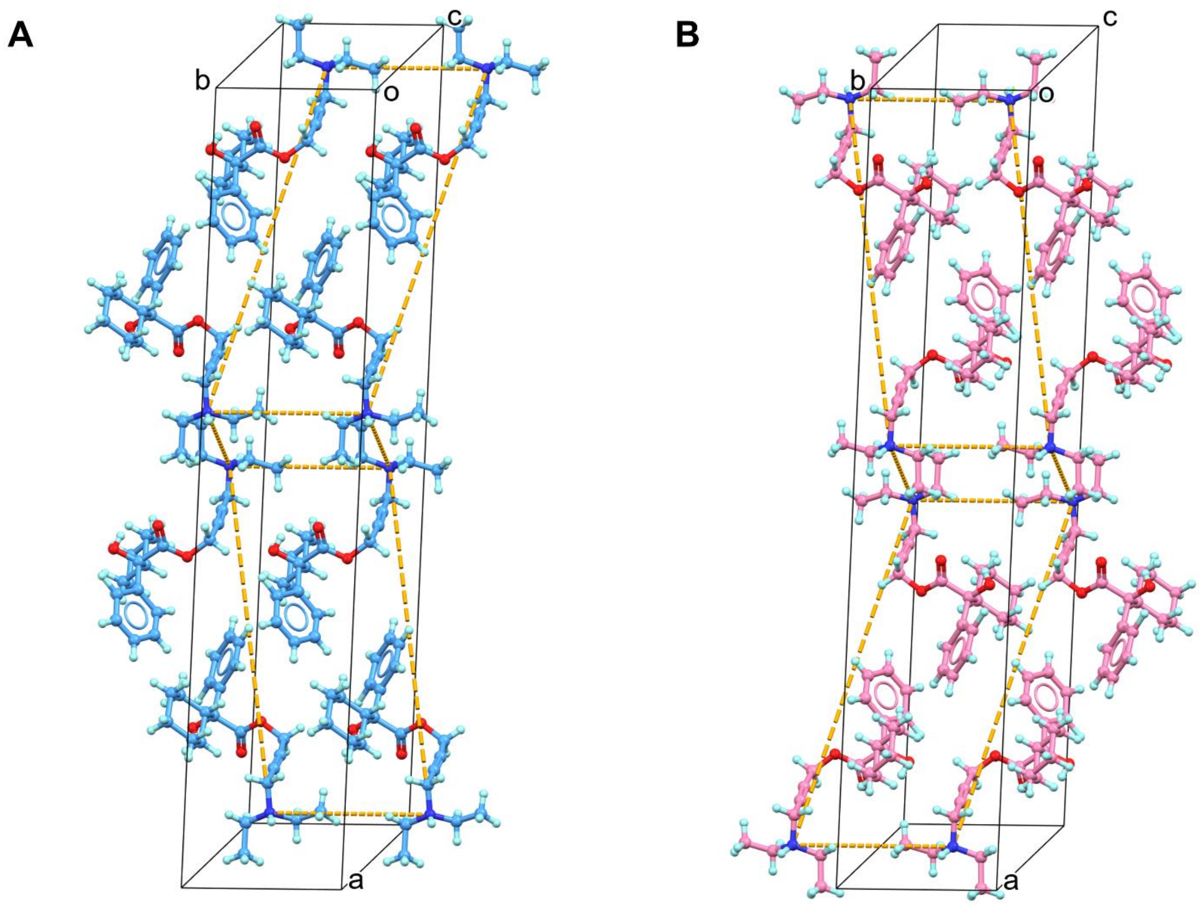


**Figure S3** (A) Packing diagram of **1R** layers; (B) Packing diagram of **1S** layers. **1R** was colored in blue, **1S** was colored in violet. $Cl^{-}$ anions were omitted for clarification. The dashed lines in orange indicated the different positions **1R** and **1S** occupied in the unit cell.

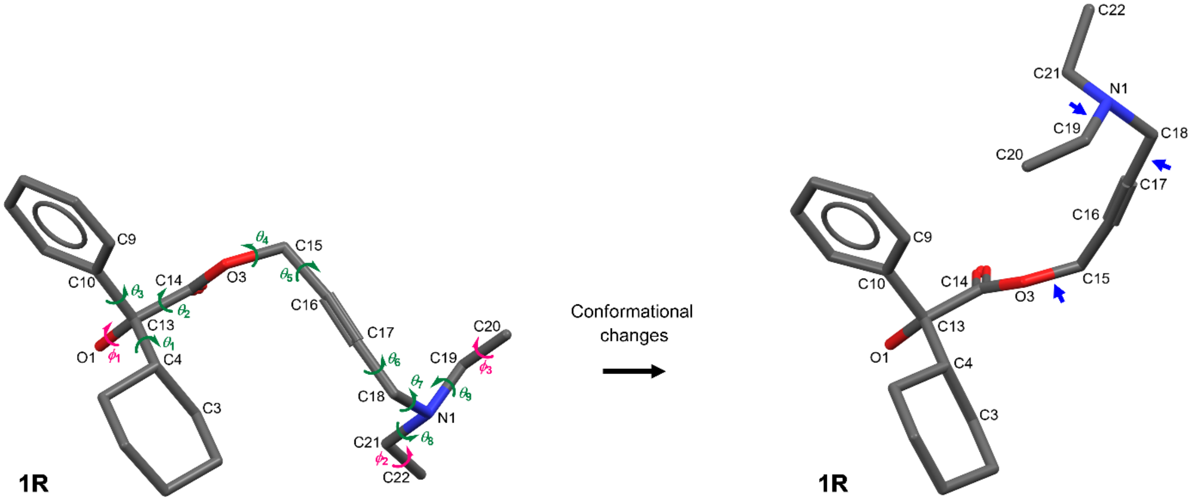


| *Drug formulation state* | *Biologically active state* |
| --- | --- |
| $\theta_1$ **(C3–C4–C13–O1)** : 61.16° | $\theta_1$ **(C3–C4–C13–O1)** : 48.81° |
| $\theta_2$ **(O3–C14–C13–O1)** : -178.89° | $\theta_2$ **(O3–C14–C13–O1)** : -157.76° |
| $\theta_3$ **(C9–C10–C13–O1)** : 162.08° | $\theta_3$ **(C9–C10–C13–O1)** : 171.7° |
| $\theta_4$ **(C14–O3–C15–C16)** : -89.25° | $\theta_4$ **(C14–O3–C15–C16)** : 140.17° |
| $\theta_5$ **(O3–C15–C16–C17)** : 25.59° | $\theta_5$ **(O3–C15–C16–C17)** : -37.00° |
| $\theta_6$ **(C16–C17–C18–N1)** : -128.98° | $\theta_6$ **(C16–C17–C18–N1)** : 144.13° |
| $\theta_7$ **(C17–C18–N1–C19)** : 56.33° | $\theta_7$ **(C17–C18–N1–C19)** : 67.58° |
| $\theta_8$ **(C18–N1–C21–C22)** : -170.32° | $\theta_8$ **(C18–N1–C21–C22)** : -92.06° |
| $\theta_9$ **(C18–N1–C19–C20)** : 58.37° | $\theta_9$ **(C18–N1–C19–C20)** : -106.85° |

**Figure S4** The conformation changes of **1R** from drug formulation state to biologically active state**.** $\theta_1$–$\theta_9$ are colored in green which dramatically influence the whole structure; $\phi_1$–$\phi_3$ are colored in red that have less impact on the overall structure. The major changes were marked with blue arrows.

**Table S1** MicroED data statistics of oxybutynin hydrochloride **1**.

| | |
|---|---|
| Stoichiometric formula | $C_{22}H_{32}ClNO_3$ |
| Molar mass | 393.94 |
| Temperature (K) | 80 |
| Crystal system | Monoclinic |
| Space group | C 2/c |
| Unit cell lengths (Å) | |
| a | 38.85 |
| b | 7.92 |
| c | 14.43 |
| Unit cell angles (°) | |
| α | 90.000 |
| β | 110.560 |
| γ | 90.000 |
| Cell volume (Å$^3$) | 4157.19 |
| No. of observed reflections | 9001 |
| No. of unique reflections | 3004 |
| $R_{obs}$ (%) | 15.9 |
| $R_{meas}$ (%) | 19.4 |
| I/Sigma | 4.50 |
| $CC_{1/2}$ | 98.8 |
| **Resolution (Å)** | **0.87** |
| **Completeness (%)** | **83.9** |
| **$R_1$ (%)** | **18.42** |
| $wR_2$ (%) | 45.40 |
| GooF | 1.442 |

**Table S2** Hydrogen bonds statistics of oxybutynin hydrochloride **1** (Å, °).

| **1R** | **D–H** | **H···A** | **D···A** | **D–H···A** |
|---|---|---|---|---|
| N1–H···Cl1 | 1.139 | 1.959 | 3.049 | 158.78 |
| O1–H···Cl1′ | 0.733 | 2.590 | 3.085 | 126.69 |
| C19–H···Cl1′ | 0.972 | 2.714 | 3.592 | 150.51 |
| **1S** | **D–H** | **H···A** | **D···A** | **D–H···A** |
| N1′–H···Cl1′ | 1.139 | 1.959 | 3.049 | 158.78 |
| O1′–H···Cl1 | 0.733 | 2.590 | 3.085 | 126.69 |
| C19′–H···Cl1 | 0.972 | 2.714 | 3.592 | 150.51 |

**Table S3** Protein-ligand interactions in $M_3R$/**1R** complex predicted by molecular docking (Å, °).

**Salt Bridges**

| Residue | AA | Distance | Protein positive? | Ligand Group[a] |
|---|---|---|---|---|
| 147 | ASP | 3.59 | No | DEA |

**Hydrogen Bonds**

| Residue | AA | Ligand Group[a] | H–A | D–A | D–H···A |
|---|---|---|---|---|---|
| 151 | SER | DEA | 3.20 | 4.10 | 157.61 |
| 507 | ASN | Carb | 2.05 | 2.94 | 144.19 |

**π-stacking**

| Residue | AA | Ligand Group[a] | Distance[a] | Angle[b] | Stacking Type |
|---|---|---|---|---|---|
| 503 | TRP | Ph | 4.91 | 69.85 | T-shaped |

**Hydrophobic Interactions**

| Residue | AA | Ligand Group[a] | Distance |
|---|---|---|---|
| 148 | TYR | DEA | 3.59 |
| 148 | TYR | Cy | 3.84 |
| 152 | ASN | Ph | 3.53 |
| 155 | VAL | Ph | 3.68 |
| 199 | TRP | Cy | 3.49 |
| 231 | THR | Cy | 3.65 |
| 234 | THR | Cy | 3.79 |
| 235 | ALA | Cy | 3.95 |
| 238 | ALA | Ph | 3.68 |
| 503 | TRP | DEA | 3.71 |
| 506 | TYR | ALip | 3.77 |
| 529 | TYR | ALip | 3.88 |
| 533 | TYR | DEA | 3.52 |

**Notes**: [a]Ligand group was represented by abbreviation: diethylamine (DEA), carbonyl group (Carb), phenyl group (Ph), cyclohexyl group (Cy), aliphatic chain (ALip); [b]Distances were measured between centroids determined as the center of aromatic rings. [c]Angles were measured by the angles between planes of two aromatic rings.

**Table S4** Protein-ligand interactions in $M_3R$/**2** complex predicted by molecular docking (Å, °).

| **Salt Bridges** | | | | | |
|---|---|---|---|---|---|
| **Residue** | **AA** | **Distance** | **Protein positive?** | **Ligand Group[a]** | |
| 147 | ASP | 3.42 | No | DMA | |
| **Hydrogen Bonds** | | | | | |
| **Residue** | **AA** | **Ligand Group[a]** | **H–A** | **D–A** | **D–H···A** |
| 151 | SER | DMA | 3.08 | 3.97 | 156.92 |
| 507 | ASN | Carb | 1.99 | 2.89 | 145.07 |
| **π-stacking** | | | | | |
| **Residue** | **AA** | **Ligand Group[a]** | **Distance[b]** | **Angle[c]** | **Stacking Type** |
| 503 | TRP | Ph | 4.81 | 62.68 | T-shaped |
| **Hydrophobic Interactions** | | | | | |
| **Residue** | **AA** | **Ligand Group[a]** | **Distance** | | |
| 148 | TYR | Cy | 3.61 | | |
| 152 | ASN | Ph | 3.57 | | |
| 155 | VAL | Ph | 3.77 | | |
| 199 | TRP | Cy | 3.54 | | |
| 231 | THR | Cy | 3.63 | | |
| 238 | ALA | Ph | 3.74 | | |
| 506 | TYR | Cy | 3.82 | | |
| 529 | TYR | ALip | 3.73 | | |

**Notes**: [a]Ligand group was represented by abbreviation: dimethylamine (DMA), carbonyl group (Carb), phenyl group (Ph), cyclohexyl group (Cy), aliphatic chain (ALip); [b]Distances were measured between centroids determined as the center of aromatic rings. [c]Angles were measured by the angles between planes of two aromatic rings.

**Table S5** Protein-ligand interactions in $M_3R$/**3** complex (PDB entry: 4U15) determined by X-ray diffraction (Å, °).

| Salt Bridges | | | | |
|---|---|---|---|---|
| **Residue** | **AA** | **Distance** | **Protein positive?** | **Ligand Group[a]** |
| 147 | ASP | 4.49 | No | BR |

| Hydrogen Bonds | | | | | |
|---|---|---|---|---|---|
| **Residue** | **AA** | **Ligand Group[a]** | **H–A** | **D–A** | **D–H···A** |
| 151 | SER | BR | 1.70 | 2.49 | 137.96 |
| 507 | ASN | Carb | 2.15 | 2.91 | 129.69 |

| π-cation Interactions | | | |
|---|---|---|---|
| **Residue** | **AA** | **Ligand Group[a]** | **Distance[b]** |
| 503 | TRP | BR | 5.57 |
| 506 | TYR | BR | 5.90 |
| 529 | TYR | BR | 4.12 |

| Hydrophobic Interactions | | | |
|---|---|---|---|
| **Residue** | **AA** | **Ligand Group[a]** | **Distance** |
| 148 | TYR | Th | 3.76 |
| 199 | TRP | Th | 3.87 |
| 225 | LEU | Th | 3.84 |
| 503 | TRP | BR | 3.76 |
| 506 | TYR | BR | 3.62 |
| 529 | TYR | BR | 3.80 |

**Notes**: [a]Ligand group was represented by abbreviation: bicyclic ring (BR), carbonyl group (Carb), thiophenyl group (Th). [b]Distances were measured between centroids determined as the center of aromatic rings.

**Table S6** Protein-ligand interactions in $M_3R$/**4** complex predicted by molecular docking (Å, °).

| Salt Bridges | | | | |
|---|---|---|---|---|
| **Residue** | **AA** | **Distance** | **Protein positive?** | **Ligand Group[a]** |
| 147 | ASP | 4.66 | No | BR |

| Hydrogen Bonds | | | | | |
|---|---|---|---|---|---|
| **Residue** | **AA** | **Ligand Group[a]** | **H–A** | **D–A** | **D–H···A** |
| 507 | ASN | Carb | 2.67 | 3.30 | 119.65 |

| π-cation Interactions | | | |
|---|---|---|---|
| **Residue** | **AA** | **Ligand Group[a]** | **Distance[b]** |
| 148 | TYR | BR | 5.38 |
| 503 | TRP | BR | 4.91 |

| Hydrophobic Interactions | | | |
|---|---|---|---|
| **Residue** | **AA** | **Ligand Group[a]** | **Distance** |
| 152 | ASN | Ph | 3.65 |
| 155 | VAL | Ph | 3.98 |
| 199 | TRP | Ph | 3.42 |
| 225 | LEU | Ph | 3.33 |
| 231 | THR | Ph | 3.36 |
| 234 | THR | Ph | 3.30 |
| 235 | ALA | Ph | 3.98 |
| 238 | ALA | Ph | 3.81 |
| 503 | TRP | BR | 3.63 |
| 503 | TRP | Ph | 3.70 |
| 529 | TYR | BR | 3.48 |
| 533 | TYR | BR | 3.46 |
| 533 | TYR | BR | 3.15 |

**Notes**: [a]Ligand group was represented by abbreviation: bicyclic ring (BR), carbonyl group (Carb), phenyl group (Ph). [b]Distances were measured between centroids determined as the center of aromatic rings.
